# ESQmodel: biologically informed evaluation of 2-D cell segmentation quality in multiplexed tissue images

**DOI:** 10.1101/2023.07.06.547438

**Authors:** Eric Lee, Dongkyu Lee, Wayne Fan, Andrew Lytle, Yuxiang Fu, IMAXT Consortium, David W. Scott, Christian Steidl, Samuel Aparicio, Andrew Roth

## Abstract

**Motivation:** Single cell segmentation is critical in the processing of spatial omics data to accurately perform cell type identification and analyze spatial expression patterns. Segmentation methods often rely on semi-supervised annotation or labeled training data which are highly dependent on user expertise. To ensure the quality of segmentation, current evaluation strategies quantify accuracy by assessing cellular masks or through iterative inspection by pathologists. While these strategies each address either the statistical or biological aspects of segmentation, there lacks an unified approach to evaluating segmentation accuracy.

**Results:** In this paper, we present ESQmodel, a Bayesian probabilistic method to evaluate single cell segmentation using expression data. By using the extracted cellular data from segmentation and a prior belief of cellular composition as input, ESQmodel computes per cell entropy to assess segmentation quality by how consistent cellular expression profiles match with cell type expectations.

**Availability and implementation:** Source code is available on Github at: https://github.com/Roth-Lab/ESQmodel under the MIT license.

## 1 Introduction

Recent advances in multiplexed spatial proteomic imaging platforms have broken new ground for large-scale studies of tumour heterogeneity. Emerging high-throughput platforms, such as imaging mass cytometry (IMC) [1], are capable of extracting signals in the form of images from around fifty protein markers from intact specimens. This enables spatially aware phenotyping of single cells as well as the visualization of protein distribution to analyze spatial organization within tissues [2, 3].

A critical step in the data processing of the hyper-stacked images is single cell segmentation [4, 5]. Single cell segmentation is the task of identifying single cells using specific protein markers that label regions such as the nucleus, cytoplasm and membrane [6]. Current widely adopted segmentation methods utilize classical image processing algorithms or employ interactive supervised machine learning techniques using platforms including ImageJ [7, 8], CellProfiler [9], Ilastik [10] and QuPath [11]. More recent work has applied existing object detection based deep learning schemas by developing models, such as Cellpose [12] and StarDist [13]. However, computational methods often require manual annotation in a semi-supervised fashion or a substantial amount of labeled training data [14]. Thus, the results of such studies are highly dependent on the expertise of the annotators. A key challenge of applying any segmentation approach is the identification of common segmentation errors. These include partially segmenting cells (partial segmentation), splitting cells apart (split segmentation), and merging cells into one entity (merge segmentation). Spotting such errors is laborious as it requires expertise to iteratively examine each image.

The current canonical approach to evaluating segmentation accuracy is through manual inspection of protein expression by pathologists or computational evaluation of segmentation masks. Some available performance metrics include mean average precision and recall with a ground truth, fraction of cell mask in the foreground, and standard deviation of cell size [15, 16]. While these metrics each assess an aspect of the segmentation quality, there are discrepancies in the interpretation of each metric. In addition, there is a lack of a unified strategy to evaluate segmentation quality that integrates both the statistical properties and and biological contexts. To address this need, we propose ESQmodel, a Bayesian probabilistic approach to evaluate the single cell segmentation quality using cellular expression profiles. Rather than directly evaluating the masks, this model leverages the extracted cell by marker expression matrix from the segmentation step and a cell type by marker prior expression matrix provided by the user. The prior expression matrix represents the belief of the expected expression profile for each cell type. We fit this model using Markov Chain Monte Carlo (MCMC) methods. ESQmodel computes a cellular entropy score of a latent variable that represents a vector of percentage contribution of each cell type’s profile to the observed cellular expression. The average entropy then serves as a proxy of the segmentation performance, assuming that well segmented cells will have expression profiles with high contribution from a single cell type’s expression profile. We tested the performance of this model on simulated, breast cancer (BC) [17], classical Hodgkin lymphoma (CHL), reactive lymph node (RLN) and tonsil IMC datasets to demonstrate generality across tissue types. Our results show that ESQmodel automates quantitative assessment by providing a biologically informed metric for single cell segmentation performance.

## 2 Methods

The objective of ESQmodel (Fig. 1) is to quantitatively evaluate the quality of cell segmentation using marker expression levels across cells. This method has two parts. First, we infer the contribution of each population to the observed expression of a segmented cell in the form of a probability vector. Then, we calculate entropy of the distribution defined by this vector, in which the resulting value indicates the degree of mixing required to explain the expression of a segmented cell. Intuitively well segmented cells will only have contributions from a single cell population (low entropy), whereas poorly segmented cells will be mixture of expression profiles from different populations (high entropy). We propose that the average entropy across cells calculated using the inferred probability distributions reflects the quality of cell segmentation. This serves as a metric to not only evaluate how cells are segmented within a single image, but also to compare between images across a multi-image dataset.

**Fig. 1.**
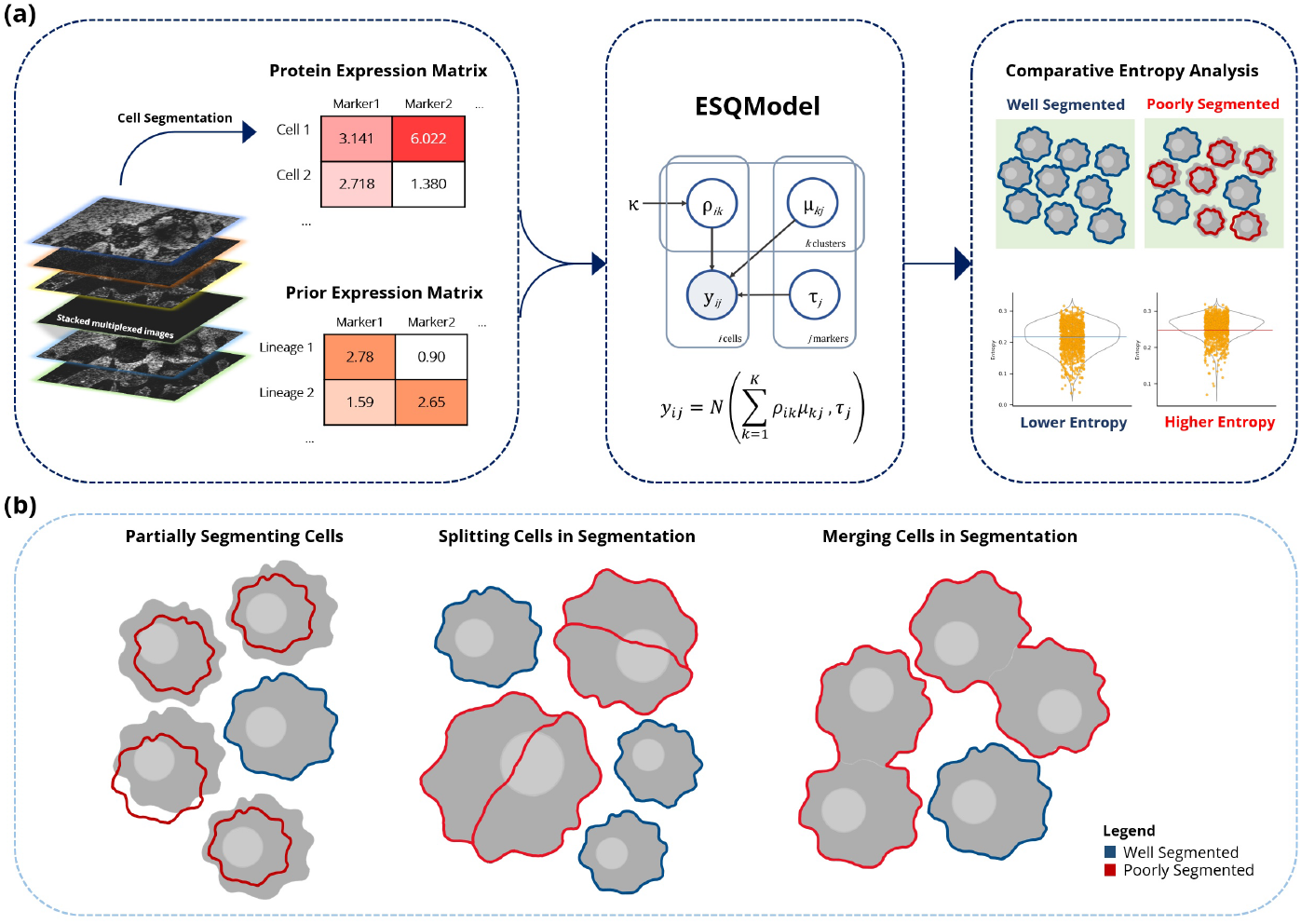
Schematic overview of ESQmodel. (**a**) ESQmodel takes in two inputs: 1) a protein expression matrix generated from applying single cell segmentation on stacked multiplexed images, 2) a user provided prior expression matrix indicating the level of cellular expression per expected cell type. ESQmodel outputs a entropy score per cell segmented. This metric can be used to compare performances of different segmentation approaches on the same dataset. (**b**) Three common errors in segmentation: partially segmenting cells, splitting cells and merging cells.

### 2.1 Model description

ESQmodel is described by the following hierarchical model:

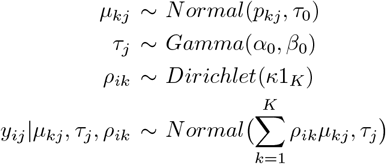

Let the measured cellular expression matrix of an image be denoted as 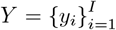, where *i* indexes cells and *I* is the total number of cells segmented in an image. The expression profile of each cell is represented by a vector of *J* measured markers which we denote by 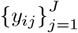, where *j* indexes markers. We assume that the expression profile of each cell is a combination of contributions from different cell types *k* ∈ {1… *K}*, where *K* is the number of cell types. This assumption reflects the fact that cell segmentation may erroneously assign regions from multiple real cells to same predicted cell mask. As such, we can associate each cell *i* with a multivariate latent vector denoted as 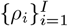 which follows a probability distribution on the simplex given by a symmetric Dirichlet distribution. Here, *ρ*_*ik*_ represents the percentage of contribution of each cell type *k* to the expression *y*_*i*_. We scale the concentration parameter of the Dirichlet by a scaling parameter *κ* defined as 0.1. This choice of hyper-parameter should encourage sparsity and prefer for *ρ*_*i*_ to assign most of the mass to a few components. This prior reflects the assumption that the segmentation masks are largely correct.

For each cell type *k*, there is an associated parameter *μ*_*kj*_ representing the mean of the expression level of marker *j* in population *k*. We assume *μ*_*kj*_ follows a Gaussian distribution, with its mean, *p*_*kj*_, set according to the input prior expression matrix *P* and precision set to a moderate value. Specifically, the prior expression matrix is a cell type by marker matrix which represents the prior belief of the marker intensity for each cell type. A method to construct prior beliefs is described previously [3]. For each measured marker, we define a precision *τ*_*j*_ which follows a Gamma prior distribution.Given the parameters we model the observed cellular expression of marker *j* in cell *i, y*_*ij*_, as a Normally distributed with mean 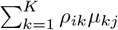 and precision of *τ*_*j*_.

### 2.2 Inference of latent variables

Due to the complexity of the posterior distribution, exact inference of the latent variables is not tractable. Instead, we implemented a Gibbs sampler to perform MCMC sampling to approximate the posterior distribution. The sampler infers the latent variables *μ, τ*, and *ρ* using the Metropolis-Hastings algorithm in an iterative fashion. All variables are randomly initialized with only *ρ* having a constraint of each vector summing to 1. The default setting of the number of iterations is set to 10000. Point estimates derived from the approximate posterior distributions for each latent variable are constructed by averaging values across iterations after removing a burn-in of the first half of the MCMC trace. Traces of each variable are plotted as an output figure to assess convergence.

### 2.3 Entropy analysis

To evaluate the segmentation quality, we compute the entropy of the point estimates of *ρ* across all cells of an image acquisition. Here, entropy represents the measure of the uncertainty of the generative process of the expression profile of a cell. A well segmented image will have entropies close to 0 which indicates a stronger contribution of singular cell type to the expression profile of cells. The overall segmentation quality can be evaluated by taking the average of the model output: a distribution of entropy across all cells.

### 2.4 Datasets

#### 2.4.1 Simulated datasets

Simulated datasets were generated by forward simulation to assess model performance against three different types of segmentation errors. First, we sampled from our described model. The expression and prior matrix was created by referencing to a labeled human bone marrow mass cytometry dataset [18]. The simulated expression were sampled from Gaussian distributions with their means set to the mean expression profiles of different cell types in the reference dataset. An arcsinh transformation was applied as normalization. Next, we introduced the errors to create each dataset: (1) we built partial segmentation datasets by sampling noise percentage variables to introduce random loss of marker intensity. The noise percentage is applied to 20% and 50% of cells respectively to create two separate datasets; (2) we built split segmentation datasets by sampling split percentage variables to divide 20% of cells apart. This splits single cells to pairs of cells each with a portion of the original total expression; (3) we built merge segmentation datasets by randomly merging 20% of cells. This combined the expression between cells and reduced the total cell count. Using this method, we simulated 45 13-marker expression matrices of 2000 cells with a cell type count enumerated from 2 to 10 each with 5 replicates. For each expression matrix, we introduced the three segmentation errors which makes a total of 180 datasets.

#### 2.4.2 METABRIC IMC dataset

36 IMC images from a published breast cancer imaging mass cytometry dataset of the METABRIC cohort [17] were analyzed to evaluate the performance on real data. Raw IMC images and the expression matrix of segmented hyper-stacked images were used. In-house segmentation was performed by using CellProfiler and Ilastik under a modified procedure of the ImcSegmentationPipeline [19]. Four datasets featuring the well segmented and the three types of segmentation errors—partial, split, and merge—were made through manual labeling using Ilastik. We subsetted the dataset by retaining only selected well-stained immune, stromal, and epithelial markers including CD45, SMA, CK5, and CK7. The original labeled cell types were also reduced to one of the seven labeled cell types: stromal, immune, CK5+, CK7+, SMA+, epithelial, and CK5+ SMA+. A prior expression matrix was generated from the original expression matrix for the seven merged cell types. All generated expression matrices were normalized by an arcsinh transformation with a co-factor of 0.8 and clipped at the 99.5 percentile per marker prior to performing inference.

#### 2.4.3 CHL, RLN, and Tonsil IMC datasets

3 IMC images each of CHL, RLN, and tonsil were generated in-house to compare segmentation accuracy on different tissue types. Procedures for IMC including antibody titration, staining, and acquisition are as previously described [17]. Three representative segmentation methods were performed on the images, including the modified procedure of the ImcSegmentationPipeline, the watershed algorithm and StarDist using QuPath. A detailed description of the procedure and settings for segmentation is described in the Supplementary Methods. Subsetting of the different datasets were done by retaining only selected lineage protein markers for representative cell types. Prior expression matrices and normalization was performed as described in the METABRIC IMC dataset section. An additionally layer of scaling was done on these datasets to assist in performance.

## 3 Results

### 3.1 ESQmodel identifies poor segmentation in simulated datasets

We first assessed the ability of ESQmodel to detect erroneous segmentation in simulated datasets. We evaluated its ability to spot three common segmentation errors: partial, split, and merge. We first generated datasets from our model with the source of average expression profiles being from an annotated human bone marrow mononuclear CyTOF dataset [18]. To simulate the impact of segmentation errors on the expression datasets, we generated 180 datasets that introduced changes in intensity and the total cell count.

The average entropy score outputted from ESQmodel reflects the uncertainty of a cell belonging to a specific cell type, which is a proxy measurement of assessing quality of segmentation. Fitting on simulated datasets, ESQmodel produced a higher average entropy score in all datasets impacted from different types of erroneous segmentation compared to the well segmented dataset (Fig. 2). For the partial segmentation dataset, we observed an average of 13% increase in average entropy when 20% of cell expressions were affected, while having a close to two fold increase after an additional 30% of the cells were reduced in expression intensity. A similar trend was observed in the split segmentation dataset, where the average entropy increased by 29.5% and 47% when 20% and 50% of the single cell expressions were split into halves. In the merge segmentation dataset, there was a 17% increase in average entropy in both percentage cases. With increases in average entropies when erroneous segmentation is present, this suggests ESQmodel is capable of capturing segmentation quality from biological expression matrices.

**Fig. 2.**
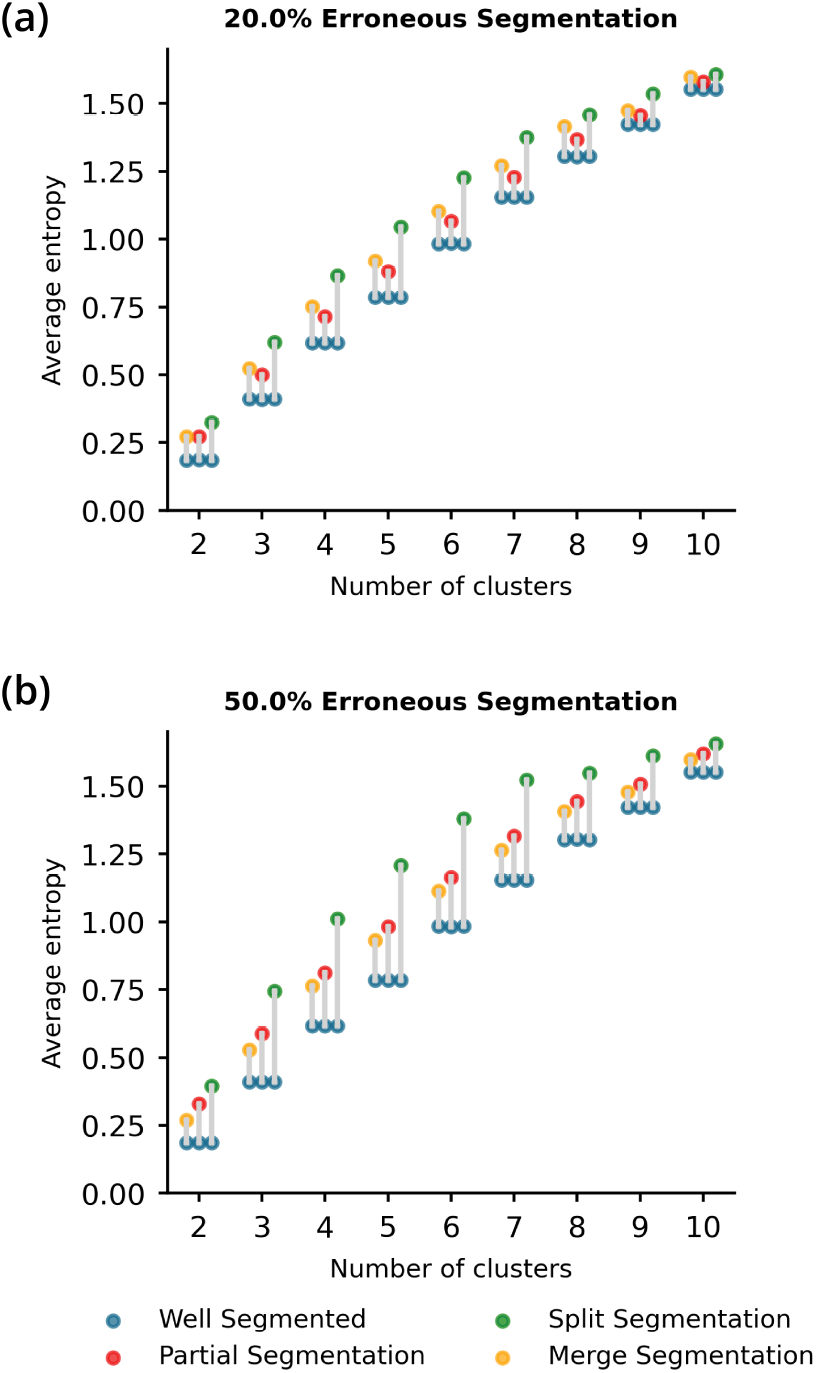
Performance on simulated data. Dumbbell plots of change in average entropy between well segmented and three cases of erroneously segmented expression data are shown. Two sets of data were created with (**a**) 20% and (**b**) 50% of total amount of cells being erroneously segmented. Colors represent different segmentation datasets.

### 3.2 Identifying introduced segmentation errors in METABRIC IMC images

We next exemplified the utility of ESQmodel on real world data by fitting with IMC images of 36 patients from the METABRIC cohort [17]. Each patient’s IMC images were manually segmented using Ilastik four times to simulate one well and three poor segmented analysis. Here we define “well” segmented as the expert performed segmentation, while “merge”, “partial”, and “split” refers to cases of segmentation that were purposely introduced respective type error.

We first evaluated ESQmodel’s ability to compare segmentation quality by using average entropy as a measure. We applied the Wilcoxon rank-sum test to determine if there is a significant difference between the set of average entropies of 36 IMC samples under well versus error introduced conditions. In all three pairwise comparisons, we observed an elevated average entropy in the error introduced segmentation compared to the well segmented (Wilcoxon, P *<* 1e-7) (Fig. 3a). This demonstrates that entropy from ESQmodel serves as an effective metric to detect the presence of erroneous segmentation.

**Fig. 3.**
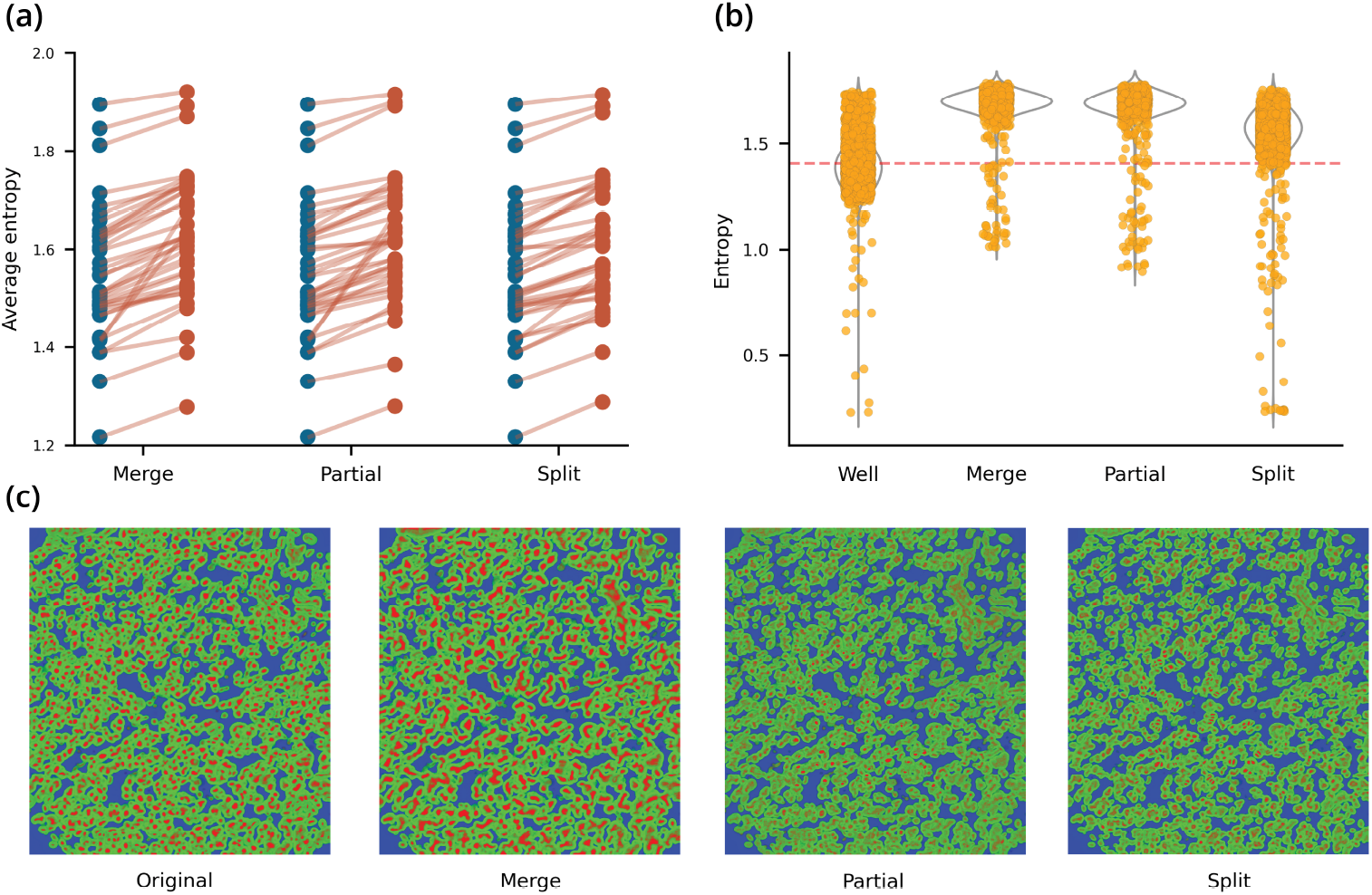
Performance on error-introduced METABRIC IMC images. (**a**) Slope chart of average entropy between well segmented and three cases of erroneously segmented expression data are shown. Blue indicates well segmentation and red indicates error-introduced. (**b**) Jitter overlayed violin plots of cellular entropy across different segmentation conditions for a single representative sample. Yellow dots are single cell entropies. Red line indicates the average entropy of the well segmented. (**c**) Ilastik probability maps of different segmentation conditions corresponding to (**b**). Red, green and yellow represent nucleus, cytoplasm and background respectively.

We then inspected the distribution and probability maps of single cell entropies in singular IMC samples. While all three erroneous cases exhibited higher average entropy than the well segmented equivalent, the entropy distribution showed distinct properties (Fig. 3b). Merge and partial segmentation have similar distributions with a higher average entropy and smaller variance than the well segmented, while the distribution for split segmentation resembled well segmented images. The probability maps of each case represented predicted cellular boundaries showing distinct qualitative trends for each type of segmentation error introduced (Fig. 3c). Merge is expected to introduce higher entropy due to drastic changes in cell expression profiles when neighbouring cells are misclassified as a single cell. As such, new cellular profiles can deviate from the prior expression matrix, thereby increasing uncertainty and entropy. A higher entropy is also expected from partial as incorrect partitioning may fail to capture protein signal in important regions of cells. Similar to merge, incomplete cell expression profiles negatively impact cell lineage identification and renders higher entropy. Split segmentation can be considered as a special case of partial where a single cell is split into relatively equal halves. We observe the increase of entropy to be moderate compared to merge and partial. An explanation for the resemblance to well segmented is that expression profiles are roughly identical in the relative intensity of each lineage marker. As the prior expression matrix represents a relative expected intensity level as opposed to an exact value, this may render split to be challenging to identify. Altogether, the distribution of entropies and the probability maps of individual IMC samples show that ESQmodel is able to produce results consistent with biological insight.

### 3.3 Analyzing the properties of ESQmodel using well segmented METABRIC IMC images

We investigated whether ESQmodel can make consistent evaluations of segmentation quality performed by different annotators. We performed manual segmentation using Ilastik on the 36 IMC images and compared the results against the published segmentation by the original author. We observed a significant correlation between the average entropies of our segmentation and the published segmentation (Pearson correlation coefficient 0.978) (Supplementary Fig. 1a). This demonstrates that ESQmodel can be used to make impartial evaluations of equally well segmented images.

We further assessed whether cellular density is a lurking variable in the calculation of cellular entropies. Denser regions in images are often more challenging to annotate which can lead to more errors in segmentation. To measure density, we computed the number of cells within 8 *μ*m radius for each cell. We observed no obvious correlation between density and average entropy (Pearson correlation coefficient 0.014) (Supplementary Fig. 1b). This suggests that the calculation of average entropies from ESQmodel are not affected by cellular density.

We next explored how the co-expression of markers affects individual cellular entropies. Using the expression matrices generated from the segmentation of the original publication, we examined the relation between spatial distribution of entropies and four selected protein markers representing immune, stromal, basal, luminal epithelial, and myoepithelial cell types: CD45, SMA, CK5, and CK7. The prior expression was defined by having only four cell types where the expectation is to have an exclusively expressing lineage marker per cell type. We observed a natural partition of the spatial organization by entropy. This is especially evident in the exemplar case in Fig. 4a where the top region has higher entropy and the bottom region lower entropy. To further explore the correlation to co-expression, we clustered cells by their expression into four distinct cell types (Fig. 4b). Using the cell type labels, we plotted the spatial cell graph labeled by its cell type (Fig. 4c). A clear correlation is shown between the spatial distribution of entropy and cluster labeling where cells that co-express CD45 and CK5 strongly yield a higher entropy than those expressing mainly a single protein marker (Fig. 4d). As co-expression of lineage markers is not defined in the prior expression matrix, cells that do not fit to expectations will have an increased entropy. This exhibits the use case of ESQmodel to identify potential segmentation errors through well-defined prior matrices, as well as identify cells that deviate from our prior expectations.

**Fig. 4.**
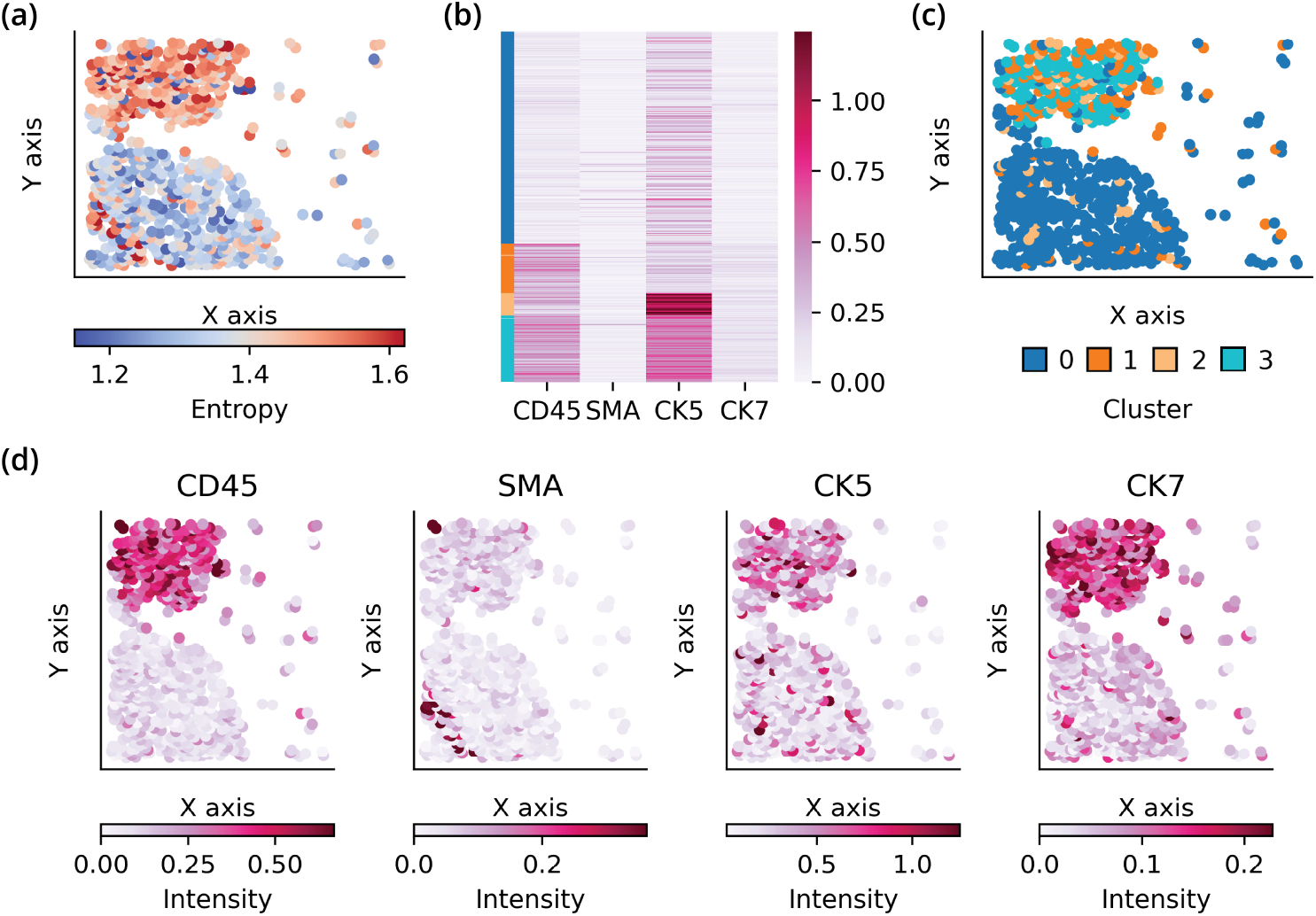
(**a**) Spatial graph of an exemplar sample. Nodes are colored by cellular entropy. (**b**) Expression heatmap with cells annotated by cell type in color.(**c**) Spatial graph colored by one of four cell types. (**d**) Spatial graph of protein markers colored by expression intensity.

### 3.4 Comparing performance of different segmentation methods on different tissue types using ESQmodel

Lastly, we compared the performance of three state-of-the-art segmentation methods on IMC images of BC, CHL, RLN and tonsil using ESQmodel. BC images were randomly subsetted from the METABRIC dataset. In-house IMC was performed on CHL, RLN, and tonsil tissues to generate image datasets. All datasets were then segmented through a semi-supervised method Ilastik, the classical image processing watershed algorithm, and a deep learning object detection method StarDist. To fully validate ESQmodel’s ability to evaluate segmentation accuracy, intentionally poor segmentation was also performed as controls by using Ilastik with erroneous manual supervision and StarDist with a heavily augmented pre-trained model. Additionally, we included and modified several other existing segmentation metrics [16] to complement our assessment on segmentation quality (Supplementary Table 1).

In Figure 5a, we illustrate segmentation performance by plotting the average entropy score for each region of interest (ROI) of the four tissue types. We first assessed the controls on Ilastik and StarDist to validate ESQmodel’s ability to serve as a comparison metric. An increase in entropy was observed in the majority of ROIs when comparing the well to poor segmentation settings. To determine if differences were significant, we applied the Student’s t test (P-value *<* 0.05) (Supplementary Table 2). For the Ilastik pair, the increases in all ROIs of RLN were statistically significant while there was a total of 4 ROIs in the other tissues that were not. We propose that as Ilastik segmentation was done in batches by tissue, some ROIs may have variable segmentation performance affected by the segmentation marks made for other ROIs. Interestingly, it was observed that poorer segmentation may identify fewer cells which are inherently easier to segment and may improve (lower) the average entropy score. For the StarDist pair, the increases in all ROIs of RLN were also statistically significant while CHL and tonsil each had one ROI that was not. For majority of the BC ROIs, StarDist in both settings did not have a significant difference in performance and was comparable to watershed. This may be due to both pre-trained models being equally unsuited for the images.

**Fig. 5.**
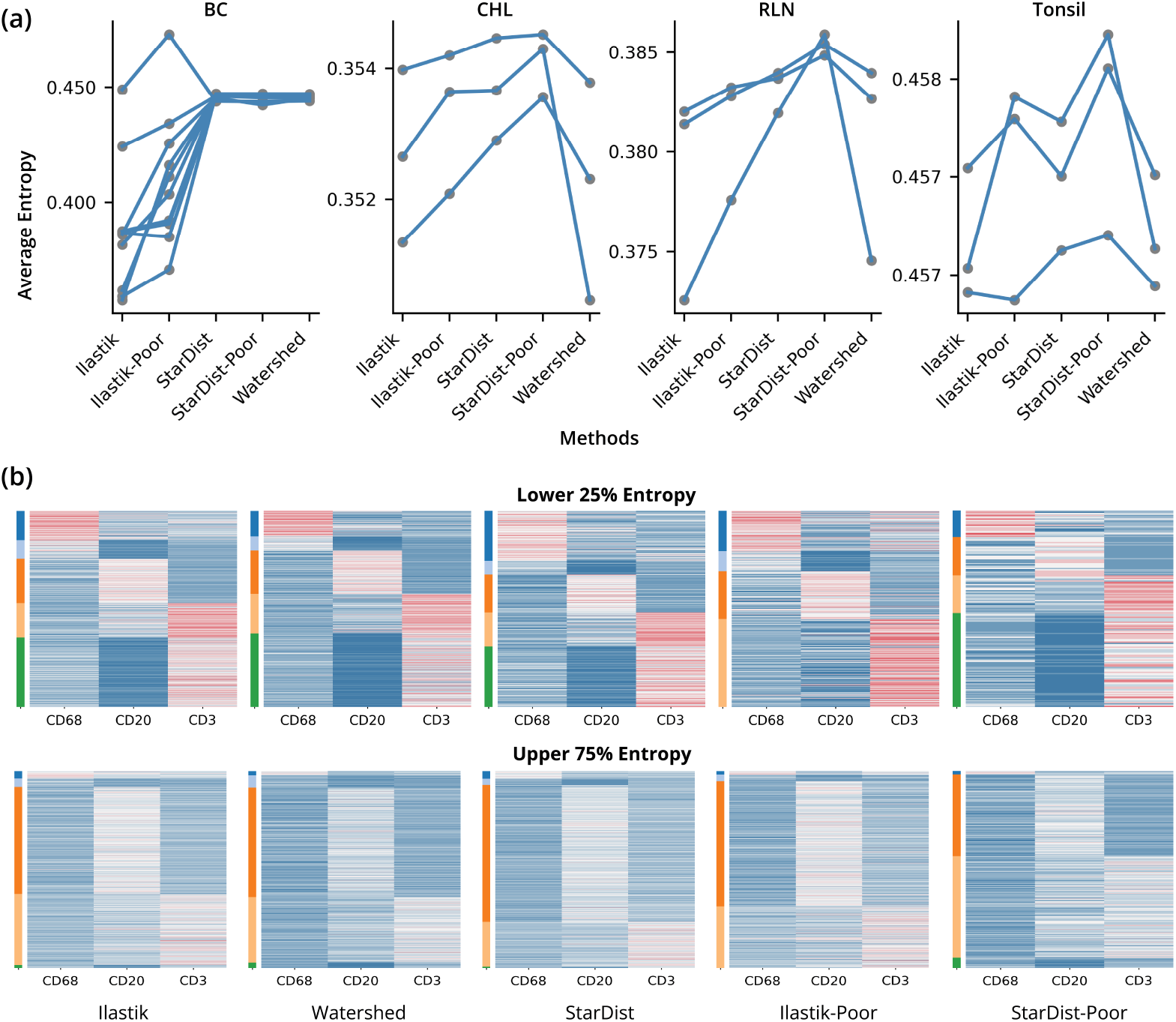
Comparison of different segmentation methods on BC, CHL, RLN and tonsil IMC experiments. (**a**) Line plots of the average entropy for each ROI of the four tissue types under five different segmentation conditions. (**b**) Expression heatmaps of the 25% of cells exhibiting lower entropy (top row) and the other 75% of cells cells exhibiting higher entropy (bottom row) for an exemplar ROI in the tonsil dataset. Heatmaps are row color-annotated by cluster labels.

We then compared the performances of each segmentation method across the four tissue types. Ilastik performed significantly better than watershed and StarDist across all ROIs in the BC and RLN datasets. There was no segmentation method that was significantly better across all ROIs in the tonsil dataset. For the CHL dataset, StarDist performed significantly worse than Ilastik and Watershed. To fully assess performance, we computed the correlation between average entropy against other existing metrics, such as a silhouette score, however there were no strong correlations (Supplementary Table 3). From the average entropy assessment, the results suggest that manual assessment with Ilastik was most effective for BC and RLN datasets, Ilastik and watershed could both be instrumental for the CHL dataset, and all methods could be considered for the tonsil dataset. The superior performance of Ilastik is consistent with expectations in which manual segmentation is often treated a gold standard against methods that are automated. Watershed can suffer from the unique properties of CHL having a heterogeneous composition of cell types with cancer cells being multinucleated. However, since cancer cells appear to be less than 1% of the tumour tissue [20], the segmentation quality of a small cell population then does not have a significant impact on the average entropy. However, as the identification of the cancer cells are vital to analysis, an assessment of the entropy and cell size for specific cells positive for CD30 may be more informative than assessing all cells as a whole. For this experiments, both StarDist models were hindered in performance as it was trained on single channel fluorescence images to fit lower resolution IMC images.

During the assessment of average entropy, we also assessed segmentation performance by visualizing expression through heatmaps. In Figure 5b, we show an exemplar ROI from the tonsil dataset. This dataset was subset to three markers: CD68, CD20 and CD3 to represent the macrophage, B cell and T cell lineage respectively. The prior expression matrix was specified with each lineage having only one marker highly expressed. We annotated cells by clustering with a variational Bayes Gaussian mixture model. Cells were then split to the well segmented lower 25% entropy and upper 75% entropy. The heatmaps for the lower entropy exhibited strong resemblance to the prior matrix while the higher entropy exhibited weaker signal across all markers for each cell. This visualization demonstrates ESQmodel’s ability to capture cells that fit with user expectation of expression intensity, and serves as a quality control metric to assess cellular expression. From the assessment, we can also observe that the three methods had similar cell type distributions agreeing with the segmentation assessment of average entropy.

To conclude, the results in Figure 5 highlight that ESQmodel is able to serve as a tool to compare performance across different segmentation tools to assist in determining a suitable choice for a dataset. In addition, ESQmodel is able to perform quality control on cell expression when having a well-defined prior expression matrix.

## 4 Discussion

In this work, we present a novel Bayesian approach, ESQmodel, that can quantitatively evaluate cell segmentation quality using expression data and prior knowledge of expected cell type composition. ESQmodel makes two important contributions to current pipelines for spatial proteomics. First, ESQmodel quantifies single cell segmentation accuracy through a biologically informed per cell entropy metric which can be used to detect the presence of erroneous segmentation. Second, ESQmodel allows for comparative entropy analysis that is instrumental in comparing performances of different segmentation methods on the same dataset. These contributions can potentially reduce the manual labor of curating segmentation by pathologists.

We emphasise that ESQmodel adds an additional metric for performing quality control of spatial single cell datasets. It is not a standalone metric however, and it should be augmented with other common metrics based on morphology and marker intensity. Indeed, there are several limitations of ESQmodel. The most significant challenge is that as the number of potential populations increases, the comparative entropy analysis becomes less effective (Supplementary Fig. 2). This is particularly apparent when designing the prior expression matrix. With more cell types, the uncertainty of cells to be identified as a cell type becomes much larger and makes entropy a less effective metric. Thus, this model is best utilized by considering a relatively small number of cell populations with distinct lineage markers. Subsequent fine grained annotation of populations can then be performed if the images pass quality control. For example, considering general T-cell lineage markers such as CD4 and CD8 for quality control using ESQmodel is preferable to inputting fine grained sub-types of T-cells that might, for example, consider different combinations of exhaustion markers. This not only increases the effectiveness of entropy as a quality control metric, but also greatly reduces running time. In addition to selecting a limited number of lineage markers, it is also vital to ensure that the markers used have high signal-to-noise ratios, since markers with a lot of non-specific staining ought to yield high entropy even with perfect segmentation.

Future work will be to establish a human-in-the-loop software to annotate high entropy cells in segmented images. This can allow trained experts to review potential erroneous annotations and make changes interactively. Another possible effort is to integrate this as a plugin for quality control of images. ESQmodel is a general model with the versatility to be applied to other spatial single cell platforms. Modifications can be made to emission distributions to suit different datasets. Additionally, we fore-see the potential for other biologically informed metrics of segmentation quality to be developed which can capture additional spurious features commonly observed in downstream analysis.

## Data Availability

Raw in-house generated datasets for all experiments used in this article have been deposited in Zenodo: https://doi.org/10.5281/zenodo.8117994. The full METABRIC dataset can be found in IDR: Image Data Resource https://idr.openmicroscopy.org/ under experimental file code: idr0076.

