## Supplementary Data for "ESQmodel: biologically informed evaluation of 2-D cell segmentation quality in multiplexed tissue images"

### Supplementary Material for Lee et al., ESQmodel: biologically informed evaluation of 2-D cell segmentation quality in multiplexed tissue images

### 1 Supplementary Note

#### 1.1 Ilastik Segmentation Settings

We performed segmentation using Ilastik [1] as part of the modified ImcSegmentation-Pipeline [2]. This modified version [3] of the pipeline allows for inputs of OME-TIFFs directly on top of MCD or TXT files which are not available for some public datasets, such as the METABRIC IMC dataset [4].

The pipeline utilizes a combination of CellProfiler [5] image processing scripts and a user created Ilastik image segmentation project. The CellProfiler scripts can be run as given with small modifications to the parameters, such as the number of markers. The procedure to setting the parameters are detailed in the README file of the repository of the modified pipeline. The Ilastik segmentation project needs to be created by the user. The procedure to segment the cells depend on the markers available in the panel. We detail the markers used for each dataset below.

- METABRIC breast cancer dataset (BC): We used Ir191 and Ir193 DNA intercalators to identify the nucleus. Cytokeratins (CK5, CK7 and panCK) were used to identify the cytoplasmic regions.
- Classical Hodgkin's lymphoma dataset (CHL): We used Ir191 and Ir193 DNA intercalators to identify the nucleus. For membrane segmentation, we used the membrane markers from the IMC cell segmentation kit (TIS-00001) from Fluidigm. CD30 was used to identify Hodgkin and Reed-Sternberg cells.
- Reactive lymph node dataset (RLN) and human tonsil dataset: We used Ir191 and Ir193 DNA intercalators to identify the nucleus. For membrane segmentation, we used the membrane markers from the IMC cell segmentation kit (TIS-00001) from Fluidigm.

To introduce the different segmentation errors in Ilastik, we used different strategies to train Ilastik to perform erroneous segmentation. For merge segmentation, we merged two nuclei together and filled in the cytoplasm that surrounds the two nucleus. For partial segmentation, we filled in only parts of the nucleus and drew the cytoplasm as a thin layer outside of the nucleus. For split segmentation, we split the nucleus into half.

#### 1.2 Watershed Segmentation Settings

We ran the watershed algorithm for cell detection in QuPath [6]. As a pre-processing step, the IMC converter [7] was used to convert MCD files to OME-TIFFs. Cell detection relies on a nuclear stain for each dataset. For all datasets, we used the Ir193 DNA intercalator. The parameters set are the following: we set the minimum area to 5 $\mu m^2$ and cell expansion was set to 2$\mu$m, intensity threshold was set to 5. The parameters were consistent across all images.

#### 1.3 StarDist Segmentation Settings

We also employed StarDist [8] for cell detection in QuPath. As a pre-processing step, the IMC converter was used to convert MCD files to OME-TIFFs. StarDist is similar to watershed as it only depends on a nuclear stain and we also used the Ir193 DNA intercalator for each dataset. For our experiments, we used single channel pre-trained models for StarDist that were developed by the StarDist developers: dsb2018_heavy_augment.pb and dsb2018_paper.pb [9]. The model that performed relatively better on our datasets was termed as the regular StarDist while we termed StartDist-Poor as StarDist used with the alternative model that had poorer performance.

#### 1.4 METABRIC IMC Subset

To demonstrate performance of ESQmodel, we used a random subset of the METABRIC IMC dataset. We list the 36 patient image codes: MB0005_1_211, MB0064_1_152, MB0109_1_487, MB0120_1_462, MB0135_1_320, MB0150_1_155, MB0199_1_157, MB0221_1_105, MB0237_1_247, MB0254_1_257, MB0272_1_285, MB0301_1_287, MB0307_1_106, MB0309_1_551, MB0312_1_178, MB0321_2_412, MB0324_1_455, MB0344_1_363, MB0354_1_36, MB0370_1_275, MB0390_1_243, MB0392_1_280, MB0394_2_417, MB0416_1_224, MB0470_1_94, MB0530_1_348, MB0549_2_547, MB0568_1_204, MB0573_1_352, MB0583_1_133, MB0594_1_498, MB0601_1_271, MB0622_1_283, MB0623_1_4, MB0646_1_506, MB0904_1_410.

For the breast cancer datasets used for comparative entropy analysis across segmentation methods, we used the following 10 images in order: MB0321_2_412, MB0005_1_211, MB0646_1_506, MB0254_1_257, MB0312_1_178, MB0120_1_462, MB0324_1_455, MB0601_1_271, MB0594_1_498, MB0392_1_280.

### 2 Supplementary Figures

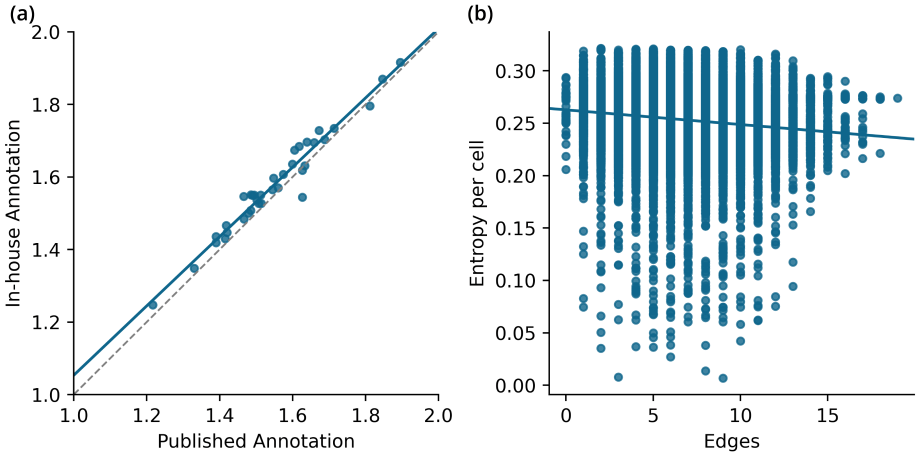


**Fig. 1** (**a**) Correlation plot between the published annotation and in-house annotation of the METABRIC IMC dataset. (**b**) Correlation plot between the number of edges per cell and cellular entropy.

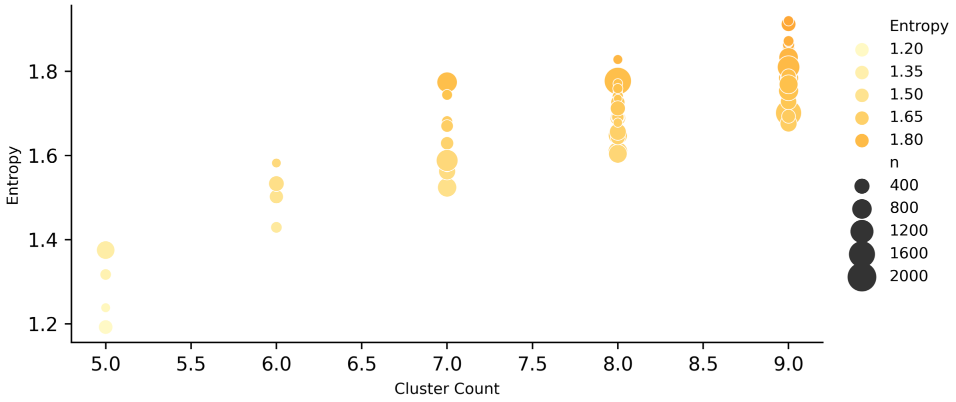


**Fig. 2** Distribution of average entropy per cluster count of the METABRIC IMC dataset. Each dot represents a segmented and processed IMC image. The cluster counts are assigned to each image as annotated in the original publication data. An increase in color gradient indicates greater average entropy. An increase in the size of dots indicates greater cell count in the images.

### 3 Supplementary Tables

#### 3.1 Supplementary Table 1. Scores of entropy from ESQmodel and various existing segmentation metrics to complement segmentation quality assessment.

| Batch | Sample | Method | Entropy | Cell Count | Cell Size Average | Cell Coverage | Silhouette Coefficient | Cell Variance | Cell Variance by Cluster |
|---|---|---|---|---|---|---|---|---|---|
| breast_1 | breast | Ilastik | 0.387472 | 1429 | 128.7845 | 0.771491 | 0.163367 | 59.09306 | 0.499279 |
| breast_1 | breast | Ilastik_bad | 0.392234 | 1536 | 63.64388 | 0.40981 | 0.224253 | 41.9341 | 0.635167 |
| breast_1 | breast | Stardist | 0.446159 | 1006 | 169.2444 | 0.713752 | 0.261732 | 113.7404 | 0.558209 |
| breast_1 | breast | Stardist_bad | 0.446485 | 761 | 189.4423 | 0.604361 | 0.315478 | 167.1526 | 0.626552 |
| breast_1 | breast | Watershed | 0.446095 | 840 | 114.5976 | 0.403543 | 0.32395 | 70.40759 | 0.568925 |
| breast_2 | breast | Ilastik | 0.448944 | 1377 | 130.4916 | 0.82446 | 0.178561 | 59.70251 | 0.458512 |
| breast_2 | breast | Ilastik_bad | 0.472962 | 1737 | 38.23316 | 0.304714 | 0.241716 | 26.43839 | 0.692228 |
| breast_2 | breast | Stardist | 0.443854 | 1199 | 122.7344 | 0.675209 | 0.289128 | 53.98037 | 0.433504 |
| breast_2 | breast | Stardist_bad | 0.443581 | 1031 | 130.9316 | 0.619379 | 0.193148 | 58.07033 | 0.423612 |
| breast_2 | breast | Watershed | 0.444 | 816 | 110.7426 | 0.414628 | 0.178977 | 68.40004 | 0.603979 |
| breast_3 | breast | Ilastik | 0.357504 | 1580 | 141.3728 | 0.82755 | 0.218895 | 64.58189 | 0.446387 |
| breast_3 | breast | Ilastik_bad | 0.416161 | 1019 | 52.48283 | 0.198136 | 0.297677 | 28.75876 | 0.578193 |
| breast_3 | breast | Stardist | 0.445972 | 1389 | 147.3575 | 0.758308 | 0.153815 | 103.8252 | 0.729873 |
| breast_3 | breast | Stardist_bad | 0.445955 | 1369 | 121.4158 | 0.615815 | 0.184279 | 60.83632 | 0.474401 |
| breast_3 | breast | Watershed | 0.446076 | 1033 | 102.4076 | 0.391926 | 0.248667 | 52.04054 | 0.487818 |
| breast_4 | breast | Ilastik | 0.386609 | 286 | 159.3636 | 0.342877 | 0.433378 | 86.64339 | 0.515929 |
| breast_4 | breast | Ilastik_bad | 0.385033 | 296 | 75.83108 | 0.168858 | 0.327287 | 50.78224 | 0.660746 |
| breast_4 | breast | Stardist | 0.446045 | 102 | 1268.06 | 0.973024 | 0.375682 | 1453.47 | 0.979966 |
| breast_4 | breast | Stardist_bad | 0.445336 | 256 | 193.9836 | 0.373584 | 0.437968 | 126.2085 | 0.546549 |
| breast_4 | breast | Watershed | 0.445204 | 218 | 109.3486 | 0.17933 | 0.406871 | 52.14638 | 0.435896 |
| breast_5 | breast | Ilastik | 0.359043 | 2159 | 116.2895 | 0.885917 | 0.477255 | 49.62916 | 0.476284 |
| breast_5 | breast | Ilastik_bad | 0.370764 | 2682 | 63.26771 | 0.598744 | 0.276843 | 38.68796 | 0.655517 |

| | | | | | | | | | |
|---|---|---|---|---|---|---|---|---|---|
| breast_5 | breast | Stardist | 0.4466 | 1376 | 178.8905 | 0.868572 | 0.480284 | 87.71788 | 0.492487 |
| breast_5 | breast | Stardist_bad | 0.446419 | 1345 | 132.439 | 0.628548 | 0.52036 | 73.50442 | 0.481552 |
| breast_5 | breast | Watershed | 0.446527 | 1227 | 113.7188 | 0.492354 | 0.480658 | 64.66152 | 0.563944 |
| breast_6 | breast | Ilastik | 0.381669 | 578 | 149.8927 | 0.599808 | 0.31848 | 71.86181 | 0.47787 |
| breast_6 | breast | Ilastik_bad | 0.403432 | 838 | 61.55728 | 0.35713 | 0.294534 | 44.12232 | 0.682423 |
| breast_6 | breast | Stardist | 0.447038 | 333 | 211.8804 | 0.488471 | 0.303746 | 228.7504 | 0.932002 |
| breast_6 | breast | Stardist_bad | 0.446905 | 447 | 127.0285 | 0.393108 | 0.307087 | 67.08503 | 0.520767 |
| breast_6 | breast | Watershed | 0.446865 | 375 | 101.704 | 0.264042 | 0.299369 | 55.10209 | 0.513809 |
| breast_7 | breast | Ilastik | 0.387198 | 1398 | 129.7876 | 0.775746 | 0.417359 | 55.95503 | 0.456529 |
| breast_7 | breast | Ilastik_bad | 0.390748 | 1339 | 97.28753 | 0.556951 | 0.428609 | 45.14269 | 0.49331 |
| breast_7 | breast | Stardist | 0.447125 | 1304 | 136.1555 | 0.759088 | 0.3564 | 57.65605 | 0.457205 |
| breast_7 | breast | Stardist_bad | 0.447126 | 1202 | 128.0237 | 0.657921 | 0.435619 | 51.56218 | 0.425416 |
| breast_7 | breast | Watershed | 0.447094 | 960 | 107.4406 | 0.44098 | 0.432804 | 59.37094 | 0.613028 |
| breast_8 | breast | Ilastik | 0.424122 | 954 | 149.4476 | 0.610915 | 0.202557 | 74.44144 | 0.491151 |
| breast_8 | breast | Ilastik_bad | 0.434159 | 1471 | 58.80421 | 0.370651 | 0.178347 | 71.51064 | 0.94782 |
| breast_8 | breast | Stardist | 0.446062 | 563 | 208.2006 | 0.502266 | 0.162354 | 238.0849 | 0.863599 |
| breast_8 | breast | Stardist_bad | 0.446045 | 716 | 143.2162 | 0.439389 | 0.157241 | 100.9656 | 0.588083 |
| breast_8 | breast | Watershed | 0.446223 | 552 | 107.7409 | 0.254838 | 0.188676 | 62.63255 | 0.570789 |
| breast_9 | breast | Ilastik | 0.36166 | 1551 | 136.9155 | 0.858129 | 0.152452 | 63.8641 | 0.496984 |
| breast_9 | breast | Ilastik_bad | 0.411047 | 1279 | 129.3839 | 0.668711 | 0.280196 | 67.21035 | 0.502806 |
| breast_9 | breast | Stardist | 0.446118 | 1104 | 190.3484 | 0.849193 | 0.173219 | 114.3841 | 0.553537 |
| breast_9 | breast | Stardist_bad | 0.44631 | 1197 | 127.8606 | 0.61847 | 0.081272 | 56.9946 | 0.447604 |
| breast_9 | breast | Watershed | 0.446361 | 1007 | 108.5472 | 0.441709 | 0.196772 | 55.2111 | 0.514959 |
| breast_10 | breast | Ilastik | 0.386081 | 1119 | 73.41912 | 0.315499 | 0.187697 | 55.6028 | 0.768593 |
| breast_10 | breast | Ilastik_bad | 0.425545 | 232 | 108.6638 | 0.096813 | 0.18849 | 138.117 | 0.861252 |
| breast_10 | breast | Stardist | 0.445228 | 137 | 119.9773 | 0.063122 | 0.143981 | 33.8707 | 0.289791 |
| breast_10 | breast | Stardist_bad | 0.442215 | 91 | 1723.886 | 0.602433 | 0.239975 | 1565.038 | 0.799028 |
| breast_10 | breast | Watershed | 0.445163 | 159 | 105.4654 | 0.064397 | 0.361927 | 36.15076 | 0.25797 |
| chl_1 | chl | Ilastik | 0.352647 | 3493 | 70.37446 | 0.983272 | 0.168073 | 30.22068 | 0.433778 |

| | | | | | | | | | |
|---|---|---|---|---|---|---|---|---|---|
| chl_1 | chl | Ilastik_bad | 0.353639 | 2027 | 108.6596 | 0.881012 | 0.16719 | 51.39988 | 0.490712 |
| chl_1 | chl | Stardist | 0.353661 | 1470 | 164.27 | 0.965908 | 0.064077 | 97.05887 | 0.614885 |
| chl_1 | chl | Stardist_bad | 0.354298 | 846 | 103.4084 | 0.349934 | 0.132801 | 57.1461 | 0.477912 |
| chl_1 | chl | Watershed | 0.350463 | 5828 | 58.52917 | 1 | 0.163498 | 34.13208 | 0.582376 |
| chl_2 | chl | Ilastik | 0.353976 | 2867 | 82.34322 | 0.944312 | 0.112449 | 35.39563 | 0.427986 |
| chl_2 | chl | Ilastik_bad | 0.354198 | 1663 | 115.6218 | 0.769116 | 0.11461 | 54.98857 | 0.480748 |
| chl_2 | chl | Stardist | 0.354461 | 1504 | 157.4126 | 0.946994 | 0.110793 | 79.89442 | 0.502144 |
| chl_2 | chl | Stardist_bad | 0.354517 | 1377 | 96.4173 | 0.531066 | 0.138971 | 34.85899 | 0.376413 |
| chl_2 | chl | Watershed | 0.353784 | 5044 | 63.36082 | 1 | 0.078362 | 34.05411 | 0.538081 |
| chl_3 | chl | Ilastik | 0.351348 | 3618 | 68.15837 | 0.986388 | 0.105101 | 27.3976 | 0.42113 |
| chl_3 | chl | Ilastik_bad | 0.352087 | 2372 | 89.59781 | 0.850104 | 0.068449 | 40.52577 | 0.458719 |
| chl_3 | chl | Stardist | 0.352896 | 1573 | 154.4054 | 0.971519 | 0.093938 | 80.91374 | 0.511746 |
| chl_3 | chl | Stardist_bad | 0.353558 | 593 | 121.6671 | 0.288594 | 0.120408 | 118.9593 | 0.692293 |
| chl_3 | chl | Watershed | 0.352312 | 3049 | 56.54444 | 0.689616 | 0.088792 | 32.49091 | 0.591423 |
| rln_1 | rln | Ilastik | 0.381391 | 2761 | 89.08874 | 0.983896 | 0.117305 | 38.53138 | 0.440933 |
| rln_1 | rln | Ilastik_bad | 0.382811 | 1841 | 130.5323 | 0.96124 | 0.18553 | 72.32596 | 0.583845 |
| rln_1 | rln | Stardist | 0.383961 | 1310 | 184.4281 | 0.966403 | 0.171286 | 97.97819 | 0.54128 |
| rln_1 | rln | Stardist_bad | 0.385403 | 835 | 115.1226 | 0.384509 | 0.128661 | 46.53942 | 0.415362 |
| rln_1 | rln | Watershed | 0.383958 | 2728 | 64.636 | 0.705308 | 0.144598 | 37.38728 | 0.579235 |
| rln_2 | rln | Ilastik | 0.382029 | 3668 | 67.13222 | 0.984964 | 0.121773 | 30.23978 | 0.450575 |
| rln_2 | rln | Ilastik_bad | 0.383229 | 2033 | 120.331 | 0.978532 | 0.1954 | 70.18375 | 0.580865 |
| rln_2 | rln | Stardist | 0.383685 | 1335 | 169.0204 | 0.902569 | 0.148343 | 95.37366 | 0.555842 |
| rln_2 | rln | Stardist_bad | 0.384855 | 195 | 224.6473 | 0.175225 | 0.166668 | 211.4131 | 0.631047 |
| rln_2 | rln | Watershed | 0.382674 | 3481 | 49.4605 | 0.688688 | 0.141525 | 32.74759 | 0.629143 |
| rln_3 | rln | Ilastik | 0.372575 | 2914 | 83.91901 | 0.97816 | 0.128813 | 37.84898 | 0.438484 |
| rln_3 | rln | Ilastik_bad | 0.377591 | 1957 | 123.0276 | 0.96306 | 0.167697 | 68.48797 | 0.571825 |
| rln_3 | rln | Stardist | 0.381952 | 1303 | 179.1235 | 0.933592 | 0.115634 | 97.30765 | 0.51091 |
| rln_3 | rln | Stardist_bad | 0.385896 | 656 | 138.8793 | 0.364419 | 0.199615 | 111.1581 | 0.730571 |
| rln_3 | rln | Watershed | 0.374561 | 2863 | 61.45721 | 0.703808 | 0.093278 | 40.57047 | 0.654407 |

| | | | | | | | | | |
|---|---|---|---|---|---|---|---|---|---|
| tonsil_1 | tonsil | Ilastik | 0.456536 | 3219 | 74.9593 | 0.965176 | 0.222476 | 35.38771 | 0.474328 |
| tonsil_1 | tonsil | Ilastik_bad | 0.45741 | 1420 | 165.3148 | 0.938988 | 0.224653 | 102.7252 | 0.547284 |
| tonsil_1 | tonsil | Stardist | 0.457283 | 1640 | 141.7276 | 0.929733 | 0.226065 | 73.68596 | 0.512099 |
| tonsil_1 | tonsil | Stardist_bad | 0.457729 | 881 | 91.42142 | 0.322169 | 0.228433 | 34.82536 | 0.366674 |
| tonsil_1 | tonsil | Watershed | 0.456635 | 2788 | 57.75323 | 0.644064 | 0.232383 | 34.96127 | 0.584197 |
| tonsil_2 | tonsil | Ilastik | 0.457046 | 3201 | 77.57013 | 0.993208 | 0.250092 | 30.49944 | 0.393793 |
| tonsil_2 | tonsil | Ilastik_bad | 0.457297 | 1586 | 153.5 | 0.973804 | 0.195729 | 97.68063 | 0.639523 |
| tonsil_2 | tonsil | Stardist | 0.457004 | 1717 | 138.6589 | 0.952309 | 0.231199 | 64.3888 | 0.458661 |
| tonsil_2 | tonsil | Stardist_bad | 0.457555 | 861 | 92.10025 | 0.317193 | 0.177994 | 38.02034 | 0.426581 |
| tonsil_2 | tonsil | Watershed | 0.457012 | 3027 | 55.82524 | 0.675932 | 0.221861 | 30.79641 | 0.547622 |
| tonsil_3 | tonsil | Ilastik | 0.456411 | 3245 | 76.79538 | 0.996804 | 0.27876 | 32.16644 | 0.420591 |
| tonsil_3 | tonsil | Ilastik_bad | 0.456373 | 1662 | 135.3051 | 0.899508 | 0.307836 | 81.94459 | 0.553778 |
| tonsil_3 | tonsil | Stardist | 0.456627 | 1643 | 143.511 | 0.943154 | 0.314531 | 65.68154 | 0.456795 |
| tonsil_3 | tonsil | Stardist_bad | 0.456705 | 644 | 102.4648 | 0.263949 | 0.21201 | 56.59264 | 0.519916 |
| tonsil_3 | tonsil | Watershed | 0.456445 | 3204 | 52.33552 | 0.670732 | 0.255183 | 28.73472 | 0.528161 |

#### 3.2 Supplementary Table 2. Pairwise comparison of segmentation performance of three representative methods as well as controls on two methods. All p-values are computed using the Student's t test. A p-value ¡ 0.05 is considered significant

| Experiment | Ilastik:Watershed | Ilastik:Stardist | Watershed:Stardist | Ilastik: Ilastik_Poor | Stardist: Stardist_Poor |
|---|---|---|---|---|---|
| breast_1 | 0 | 0 | 0.723 | 1.20E-06 | 0.069 |
| breast_2 | 3.30E-08 | 6.40E-12 | 0.321 | 2.89E-236 | 0.055 |
| breast_3 | 0 | 0 | 0.173 | 1.89E-281 | 0.821 |
| breast_4 | 6.00E-161 | 5.96E-96 | 6.59E-04 | 0.365 | 0.005 |
| breast_5 | 0 | 0 | 0.495 | 1.24E-53 | 0.123 |
| breast_6 | 1.39E-137 | 1.41E-125 | 0.513 | 4.56E-27 | 0.566 |
| breast_7 | 0 | 0 | 0.775 | 0.003 | 0.992 |
| breast_8 | 1.13E-84 | 4.50E-85 | 0.251 | 3.62E-35 | 0.905 |
| breast_9 | 0 | 0 | 0.045 | 7.69E-288 | 0.104 |
| breast_10 | 1.06E-81 | 6.36E-72 | 0.83 | 1.59E-53 | 2.60E-07 |
| chl_1 | 1.36E-55 | 4.45E-12 | 6.46E-59 | 3.18E-14 | 1.88E-04 |
| chl_2 | 0.05 | 8.29E-05 | 3.75E-08 | 0.062 | 0.675 |
| chl_3 | 5.34E-13 | 1.65E-21 | 1.20E-04 | 5.26E-07 | 0.002 |
| rln_1 | 1.11E-23 | 6.72E-15 | 0.991 | 2.33E-06 | 4.48E-07 |
| rln_2 | 9.16E-07 | 1.40E-20 | 2.93E-10 | 5.75E-15 | 4.83E-04 |
| rln_3 | 1.48E-05 | 2.18E-71 | 1.12E-47 | 2.27E-26 | 3.60E-26 |
| tonsil_1 | 0.357 | 1.32E-09 | 2.74E-07 | 1.20E-11 | 0.005 |
| tonsil_2 | 0.769 | 0.764 | 0.958 | 0.072 | 0.004 |
| tonsil_3 | 0.797 | 0.172 | 0.249 | 0.805 | 0.746 |

#### 3.3 Supplementary Table 3. Comparison of various existing segmentation metrics with entropy from ESQmodel. Spearman's rank correlation coefficient was calculated for each pair of metrics across tissue types.

| Sample | Cell Count Correlation | Cell Size Average Correlation | Pixel Coverage Correlation | Cell Coverage Correlation | Cluster Homogeneity Correlation | Cell Variance Correlation |
|---|---|---|---|---|---|---|
| breast | 0.327611307 | 0.318722476 | 0.255424174 | 0.989173581 | 0.456163798 | 0.220482979 |
| chl | 0.016381307 | 0.061951007 | 0.016381307 | 0.107693038 | 0.703901002 | 0.139595129 |
| rln | 0.003921191 | 0.0894859 | 0.003921191 | 0.036521823 | 0.03506666 | 0.051725498 |
| tonsil | 0.008230371 | 0.177342147 | 0.008230371 | 0.420104687 | 0.011457542 | 0.482898756 |
